# Prevalence of *Chlamydia trachomatis* and *Candida albicans*; hospital based study

**DOI:** 10.1101/701201

**Authors:** Albert Njeru, Joseph Mwafaida

## Abstract

**Background:** Chlamydia and candidiasis have only mild or no symptoms at all. When symptoms develop, they are often mistaken for something else like urinary tract infections or yeast infections. These infections affect both men and woman of all backgrounds and economic levels.

**Objective:** The prevalence of *Chlamydia trachomatis* (*C.trachomatis*) and *Candida albicans* (*C.albicans*) infections among attendees of Kilifi Medical Centre in Kilifi county, Kenya was studied.

**Methodology:** Urethral and vaginal swabs samples were aseptically collected from 305 subjects, examined for *Candida albicans* and *Chlamydia trachomatis* using standard microbiological methods. The swabs were analyzed using direct wet smears, Gram-stained smear and culture techniques.

**Results:** Of the 305 participants, 181 (59.34%) females and 124 (40.66%) males were tested with the overall prevalence of 53.44 % for both *Chlamydia trachomatis* and *Candida albicans* with females having a higher infection rate (35.14 %) for chlamydia and candidasis than men (17.71). Amongst the different age groups investigated, candida and chlamydia distribution was highest in participants aged 28-32 years (21.97 %).The infection rate of *C.trachomatis* (14.43 %) among the male participants was higher than the infection rate revealed among the female participants of 1.97 % while the infection rate of *C.albicans* was higher among the female participants (33.77 %) compared to the 3.28 % recorded in male participants with no co-infections revealed.

**Conclusion:** The results of this study demonstrated a significant difference between male and female chlamydia and candida infections with women being severely affected than men. The study recommended routine screening for sexually transmitted infections (STIs) which is essential in preventing infections transmissions, assessment of the role of socio-demographic and behavioral risks on *Chlamydia trachomatis* and *Candida albicans*, proper treatment of all candida and chlamydia by use of correct/effective medicines, contact tracing and treatment of sexual partners and health education.

## Introduction

Sexually transmitted infections (STIs) refers to a group of infectious (communicable) diseases which are transmitted through sexual contact and are among the major causes of illnesses globally especially in the developing countries ^1^. They are classified according to the type of organism causing the infection which could be bacterial, fungal, viral or parasitic origin. Some of the common sexually transmitted diseases include: Candidiasis, Bacterial vaginosis, herpes, Chlamydia, trichomoniasis, gonorrhea, Hepatitis B virus, HIV and syphilis ^2^.

Chlamydia is one the most frequently reported bacterial sexually transmitted diseases (STDs) and it is caused by *Chlamydia trachomatis* bacterium ^3^. The infection is adverse if not treated within the appropriate duration affecting the urethra and rectum for both sexes. The cervix also gets infected in females. Chlamydia infection may also affect some other parts of the body such as lungs, liver, throat and eyes ^4^.

Candidiasis, thrush, or yeast infection is a fungal infection (mycosis) of any species from the genus *Candida* (one genus of yeasts).*Candida albicans* is the most common agent of candidiasis in humans. Candidiasis encompasses infections that range from superficial, such as oral thrush and vaginitis, to systemic and potentially life-threatening diseases ^5^. Candidiasis is a very common cause of vaginal irritation (vaginitis) and can also occur on the male genitals ^6^. *Chlamydia trachomatis* and *Candida albicans* are among the many related factors that affect the reproductive health. Infections by these micro-organisms lead to health and economic consequences including pains, organs damage, disabilities such as blindness, deafness, infertility, insanity, paralysis and even death ^7^.

*Chlamydia trachomatis* and *Candida albicans* infections been associated with a number of adverse pregnancy outcomes including spontaneous abortion, stillbirth, prematurity, low birth-weight, postpartum endometritis, early onset of labour including premature rupturing of membranes, cervical and other cancers, chronic hepatitis, pelvic inflammatory diseases and various sequalae in surviving neonates ^8^.Documented evidence indicates that STDs can be transmitted from a pregnant mother to the baby, before; during or after the baby’s birth. It is estimated that the number of pregnant women with STDs is increasing by about 250 million a year in the developed countries and double that number in the developing countries ^9^.

Sexually Transmitted infections (*Chlamydia trachomatis* and *Candida albicans)* are becoming a major health problem. Studies suggest that knowledge on these diseases tend to be scanty and only misconceptions are widely spread. *Chlamydia trachomatis* and *Candida albicans* cause mild infections in both male and females which may sometimes go untreated ^10^. Due to the fact that *Chlamydia trachomatis* and *Candida albicans* are mostly asymptomatic, they have potentially very serious long term effects if left untreated. Untreated infections may become chronic and have long term effects on fertility and sexual health of the victims. Infected individuals have most probably entered into unprotected sex life which is an indicator of possible more serious infections including HIV/AIDS ^11^. In Kilifi Municipality the highest population is youths aged between 18-35 years an age at a greater risk of contracting STIs ^12^.The aim of this study was to document the pattern of common STIs (*Chlamydia* and Candidiasis) and to evaluate the frequency of occurrence of *Chlamydia trachomatis* and *Candida albicans* isolates among attendees of Kilifi Medical Centre.

### Methodology

Kilifi Medical Centre is a middle level hospital located in Kilifi County of Kenya. This study focused on all the participants visiting the hospital who showed symptoms of *Chlamydia trachomatis* and *Candida albicans* and only those who were referred to the laboratory for these tests over the study period. The participants were informed that their samples were to be used for study purposes. Samples from the participants who agreed to participate, were collected by the laboratory technologist for the purpose of this study. The participants payed no other extra costs for the tests. The sample size in this study was 305 participants.

### Sample collection

Sterile swabs were used to collect two samples from each tested individual. Urethral swabs and vaginal swabs were collected from males and females respectively and subjected to direct examination by wet preparation. The swabs were smeared on CLADE media and cultured for 24 hours. Gram stain was carried out on both swabs and examined with x100 objective lens under oil immersion for Gram negative diplococci and clue cells.

## Procedure

### Isolating *Chlamydia trachomatis*

Wet mounts of all swab samples were made in sterile normal saline on clean slides and examined under the low power (10x) and high power (40x) magnifications for *Chlamydia trachomatis*. Gram stain was carried out on the swabs and examined with 100x objective under oil immersion for Gram negative diplococci and clue cells. Vaginal swab specimens were inoculated into blood agar at 25 °C for 24 hours before observation

### Isolating *Candida albicans*

Diagnosis of yeast infection was done by culture and microscopic examination. A swab of the affected area was placed on a microscope slide. A single drop of 10% potassium hydroxide (KOH) solution was then added to the specimen. The KOH dissolves the skin cells, but leaves the *Candida* cells intact permitting visualization of pseudohyphae and budding yeast cells typical of many *Candida* species. For the culturing method, the sample swabs were streaked on a culture medium and incubated at 25 °C for 48 hours before observation.

### Data analysis

All the data collected was entered into Microsoft excel spreadsheets, cleaned and analyzed using R statistical package software version 3.4.0.

## Results

The table below shows that out of the 305 participants, 181 (59.34%) females and 124 (40.66%) males were tested with the overall prevalence of 53.44 % for both *Chlamydia trachomatis* and *Candida albicans*. Amongst the different age groups investigated, candida and chlamydia distribution was highest in participants aged 28-32 years (21.97 %).The infection rate of *C.trachomatis* (14.43 %) among the male participants was higher than the infection rate revealed among the female participants of 1.97 % while the infection rate of *C.albicans* was higher among the female participants (33.77 %) compared to the 3.28 % recorded in male participants no co-infections revealed.

The prevalence rate of both *C.trachomatis* and *C.albicans* among the female participants was 35.74 % with *C.albicans* infection accounting for 33.77 % against the *C.trachomatis* which accounted for a 1.97 % infection rate. Among the 40.66 male participants, *Chlamydia trachomatis* infections translated to 14.43 % infection rate which was higher than *Candida* infections (3.28 %).In male participants aged 28-32 years, *C.trachomatis* infections were high (7.54%) while no *C.trachomatis* infections were observed among the males aged 23-27 years. No *Candida albicans* infections were recorded in male participants aged between 18-22 years while 28-32 years had an infection rate of 2.30 % for *C.albicans.* In female participants, *Candida albicans* infection was relatively high in all age groups with high infections being recorded for the participants aged between 28-32 years while *Chlamydia trachomatis* infections were low with no infections recorded in participants aged 18-22 and 28-32 years as shown on the table below.

**Table 1:** Prevalence of *Chlamydia trachomatis* and *Candida albicans* among patients attending Kilifi Medical Centre.

| Age<br>( Years) | N<br>(%)=number<br>tested | n (%)=<br>prevalence<br>rate | No. of Males infected (%) |  | No. of Females infected (%) |  |
| --- | --- | --- | --- | --- | --- | --- |
|  |  |  | *C.trachomatis* | *C.albicans* | *C.trachomatis* | *C.albicans* |
| 18-22 | 56 (18.36) | 22 (7.21) | 4 (1.31) | 0 (0) | 0 (0) | 18 (5.90) |
| 23-27 | 81 (26.56) | 32 (10.49) | 0 (0) | 2 (0.66) | 4 (1.31) | 26 (8.52) |
| 28-32 | 92 (30,16) | 67 (21.97) | 23 (7.54) | 7 (2.30) | 0 (0) | 37 (12.13) |
| ≥ 33 | 76 (24.92) | 42 (13.77) | 17 (5.57) | 1 (0.33) | 2 (0.66) | 22 (7.21) |
| Total | 305 (100) | 163 (53.44) | 44 (14.43) | 10 (3.28) | 6 (1.97) | 103 (33.77) |

## Discussion

*Candida albicans* and *Chlamydia trachomatis* infections have been recognized as a major public health problem in Kenya. These infections are among the most common sexually transmitted infections in outpatient clinics representing up to 10 % of the caseload at many health facilities ^13^. The overall prevalence of these infections in the general population in Kilifi is unknown. The findings of this study were within the same limits as established by another study carried out in Mombasa, Kenya among the general population which estimated the prevalence rate of chlamydia to be 8.8 % and candida 19.9% ^14^. The findings of this study on the infection rates of were comparatively higher than demonstrated other studies like the study on male transport workers in Mombasa, Kenya which found the prevalence of chlamydia to be 3.6 % ^15^ and a population based multi-center study in Kisumu, western Kenya that revealed the prevalence rates for candida as 2.6 % and 3.4 % for chlamydia for men and prevalence rates were 0.9% chlamydia and 4.5% for candida in women ^16^.

Amongst the different age groups investigated, candida and chlamydia distribution was highest in participants aged 28-32 years. The results of this survey were in agreement with generally observed incidences of candida and chlamydia by the number of cases treated each year being highest among the persons aged 15-30 years old ^17^. Similarly, earlier data from studies by Aboyeji and Nwabuisi ^18^ reported that those in the age group of 15-30 years were the most infected (100%) by one sexually transmitted infection or the other in their separate studies. The World Health Organization (2006) concludes that these age groups (15-30years) are persons with the greatest sexual activity and that incidence decreases with age ^7^.

In this study, the peak age group of subjects positive for candida and chlamydia ranged from 28 to 32 years (21.97%), and vast majority of them were female (12.13 %), thus constituting the major bulk of the participants. This is a matter of concern in the context of HIV transmission as genital ulcer facilitates the transmission of and enhances susceptibility to HIV infection by sexual contact while non-ulcerative sexually transmitted infections like candida and chlamydia increases shedding of the HIV virus in the genital tract by recruiting HIV-infected inflammatory cells as part of normal host response ^19^. Other studies have also showed predisposing factors for the incidences of chlamydia and candida including multiple sex partners within less than 30 days and sexual intercourse during menses in the previous 6 months ^20^. Several studies have demonstrated that circumcision of men does not reduce their risk of acquiring these non-ulcerative sexually transmitted infections like chlamydia and candidiasis ^21^.

## Conclusion and recommendations

The results of this study demonstrated a significant difference between male and female chlamydia and candida infections with women being severely affected than men. The study recommended routine screening for sexually transmitted infections (STIs) which is essential in preventing infections transmissions, assessment of the role of socio-demographic and behavioral risks on *Chlamydia trachomatis* and *Candida albicans*, proper treatment of all candida and chlamydia by use of correct/effective medicines, contact tracing and treatment of sexual partners and health education.

## Ethical Statement

Prior to commencing the study, the proposal was approved by the Pwani University Ethics Review Committee (PUERC). A consent form was also filled by participants whose swab samples were used for the research. All data collected was treated with uttermost confidentiality and was only used for this research. All the infected participants were referred to the physicians for management under no extra cost.

## Acknowledgements

We would like to acknowledge the Pwani University School of Biological Sciences, Kilifi Medical Centre administration, laboratory staff and all the study participants.

